# InSpectro-Gadget: A tool for estimating neurotransmitter and neuromodulator receptor distributions for MRS voxels

**DOI:** 10.1101/2023.11.02.565296

**Authors:** Elizabeth McManus, Nils Muhlert, Niall W. Duncan

## Abstract

Magnetic resonance spectroscopy (MRS) is widely used to estimate concentrations of glutamate and γ-aminobutyric acid (GABA) in specific regions of the living human brain. As cytoarchitectural properties differ across the brain, interpreting these measurements can be assisted by having knowledge of such properties for the MRS region(s) studied. In particular, some knowledge of likely local neurotransmitter receptor patterns can potentially give insights into the mechanistic environment GABAand glutamatergic neurons are functioning in. This may be of particular utility when comparing two or more regions, given that the receptor populations may differ substantially across them. At the same time, when studying MRS data from multiple participants or timepoints, the homogeneity of the sample becomes relevant, as measurements taken from areas with different cytoarchitecture may be difficult to compare. To provide insights into the likely cytoarchitectural environment of user-defined regions-of-interest, we produced an easy to use tool InSpectro-Gadget, that interfaces with receptor mRNA expression information from the Allen Human Brain Atlas. This Python tool allows users to input masks and automatically obtain a graphical overview of the receptor population likely to be found within. This includes comparison between multiple masks or participants where relevant. The receptors and receptor subunit genes featured include GABA- and glutamatergic classes, along with a wide range of neuromodulators. The functionality of the tool is explained here and its use demonstrated through a set of example analyses. The tool is available at https://github.com/lizmcmanus/Inspectro-Gadget.

## Introduction

Magnetic resonance spectroscopy (MRS) is increasingly used to measure metabolite concentrations in the human brain *in vivo*. The technique has been applied across a variety of domains, including in psychiatric (1, 2), neurological (3–5), and psychological research (6, 7). This has helped drive an increased understanding of multiple facets of brain metabolic and signalling processes (8, 9). Of particular interest to many brain scientists are measurements of glutamate and γ-aminobutyric acid (GABA). These are respectively the primary excitatory and inhibitory neurotransmitters in the human brain, and so their study can potentially elucidate a wide range of questions about the mechanisms underlying behaviour and cognition (10). In contrast to the majority of other neurotransmitters and modulators in the brain, both of these transmitters are visible to MRS at widely available field strengths through the use of a number of optimised acquisition sequences (11–13).

Most MRS studies in humans to date have employed a single voxel approach that measures metabolites from a specific region of the brain during each acquisition. Common volumes for these voxels range from 3.3 to 27 ml, depending on the particular metabolites being targeted, field strength, and the acquisition sequence used. Relatively large volumes of brain tissue such as these contain many millions of neurons, along with blood vessels, cerebrospinal fluid, and white matter tracts. The GABA and glutamate signals obtained in an MRS scan thus originate from a variety of compartments, including neurons, astrocytes, and extracellular spaces. This mixture of signal sources within MRS measurements suggests general limitations on the inferences that can be supported by them, given the ambiguity around which particular biological feature might underlie any measurement variance (14, 15).

A potentially important aspect of this source-specificity problem for MRS is cytoarchitectural variability. The types of neurons that are present within an area, along with their relative density and proportions, can differ depending upon where in the brain is being investigated (16, 17). For example, an MRS voxel located in the occipital cortex will be sampling from a relatively large number of layer IV stellate neurons (18), whilst one in the insular cortex will include populations of layer V von Economo cells that would not be seen in other parts of the brain (19). In addition to this, we also see cytoarchitectural variation within larger scale divisions such as those mentioned. The occipital cortex, for example, has distinct cytoarchitectural patterns across its functional subdivisions (V1, V2, etc.) (20).

An element of this cytoarchitectural variation is differences in the specific GABA and glutamate receptors present on different cell types. In the case of glutamate, for example, neurons can express a range of ionotropic (i.e., AMPA, kainate, or NMDA) or metabotropic (i.e., mGlu) receptors, each of which differ in the form and duration of response they induce within the cell. This ionotropic/metabotropic distinction can be seen in the case of GABA receptor types also (i.e., GABA_A_ compared to GABA_B_). At the same time, one receptor type can be composed of different subunits, with different subunit combinations often affecting receptor functionality. For example, the specific subunits that comprise the tetrameric structure of the AMPA receptor determine functional properties such as the ions to which it is permeable, in turn influencing the intracellular processes induced by the receptor’s activation (21, 22). Similarly, variety in the subunits comprising GABA-A receptors is connected to differences in the timing and duration of their influence on cell polarisation (23, 24). This structural variation can therefore have an influence on the functional properties that track the cytoarchitectural composition of a particular brain area (18).

At the same time, variation in receptors for neuromodulators, such as serotonin or noradrenaline, may also represent an additional layer to interpretations of MRS estimates of glutamate and GABA concentrations. Such neuromodulators often act on families of receptor types that can vary in their effect upon cells. For example, dopamine receptors in the D_1_ family will generally stimulate cell activity whilst those in the D_2_ family will inhibit it (25), with similar response variability see for other neuromodulators too. Neuromodulator receptors are also differentially distributed across the brain in a functionally relevant manner (26–28). As such, the specific neuromodulator receptor environment in a region of interest, along with the types of GABA and glutamte receptors present, are likely to be of relevance when developing mechanistic models of that region’s function in during particular behaviours. Similarly, knowledge of the local receptor landscape may also improve interpretations of changes in neurological and psychiatric groups, given the fundamental role that neurotransmitter and neuromodulator systems are thought to play in many disorders.

One potentially powerful tool for investigating receptor distributions across the brain is the analysis of gene expression (29, 30). This approach can give relative expression levels for genes encoding specific receptor proteins, allowing receptor distributions to be studied down to the subunit level. With gene expression analysis giving information about a large number of genes simultaneously, it becomes possible to build up an expression profile for a given region that includes the full range of GABA and glutamate receptor subunits, plus the variety of neuromodulator receptors also present.

A set of gene expression data for the human brain is available through the Allen Human Brain Atlas (www.brain-map.org). This includes expression levels for many genes from several thousand sampling points across the cortex. Although this data is freely available, its use requires some degree of preprocessing and data manipulation to get information for a particular area of the brain (such as the area covered by a specific MRS voxel). This makes the use of this data somewhat technical and inconvenient, limiting the range of researchers able to integrate it with their MRS data analysis.

We therefore aimed to create an easy to use tool that interfaces with the Allen database to allow researchers to look at the different receptors likely to be found in any MRS voxel (or general region of interest) that they define. The tool created - called InSpectro-Gadget - is written in Python and is designed to require minimal user input for ease of use. This work sets out the functioning of the tool and presents some example analyses. These examples illustrate the different analysis options available to the user and demonstrate the output obtained. It is hoped that the tool will allow users to enhance their understanding and interpretation of metabolite quantities measured within an MRS voxel. In addition to interpreting already collected MRS data, the tool may also be of use when planning voxel placement where interpretation of mechanistic aspects.

The tool is available at https://github.com/lizmcmanus/Inspectro-Gadget.

## Methods

InSpectro-Gadget allows a user to enter a mask image (or images) to obtain a PDF report detailing estimates of relative expression of GABAergic, glutamatergic, dopaminergic, serotonergic, cholinergic, and adrenergic receptor subunit expression within the region specified. This information is shown in the form of both radar and violin plots to allow easy comparison between different subunits and to show the degree of variability in these measures across the region. As MRS measures of GABA and glutamate are often interpreted as representing the ratio between excitation and inhibition in a region, an estimate of this value is provided for the input region (31). The receptor genes for which expression estimates are provided are presented in Table 1.

**Table 1.**
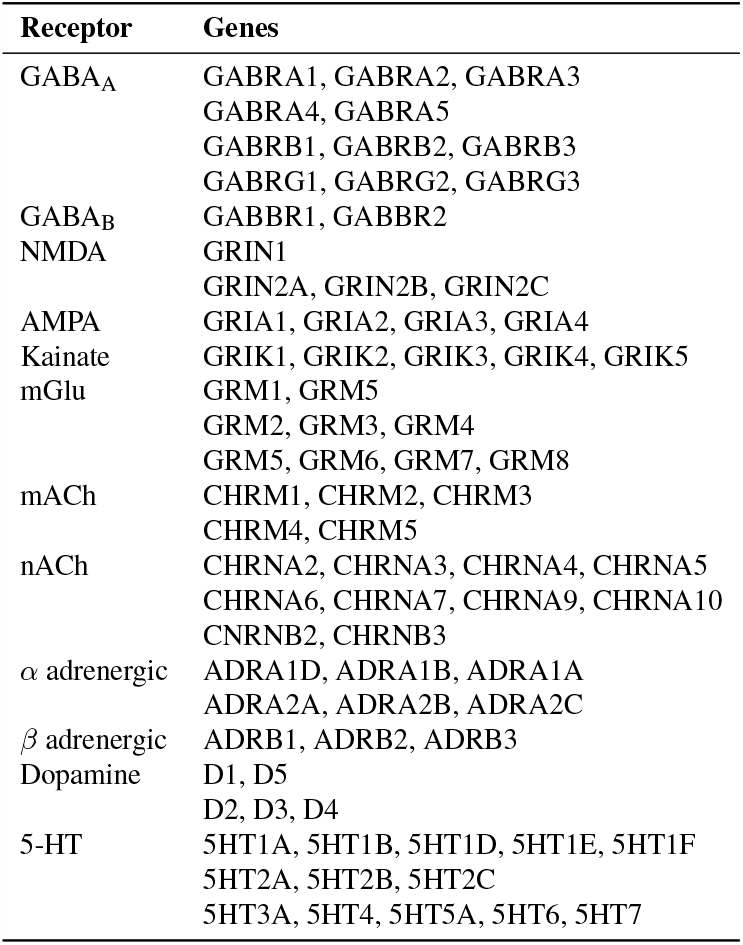
Receptor types and the specific genes related to each included in reports. mACH = muscarinic acetylcholine; nACh = nicotinic acetylcholine; 5-HT = 5-hydroxytryptamine; mGlu = metabotropic glutamate receptor

The tool provides three different analysis options. Firstly, a user can enter a single mask to obtain a gene expression report for that region. Secondly, a user can enter a pair of masks in order to compare gene expression between those regions. Finally, the user can enter a collection of masks from different participants in order to identify potential variability in receptor distributions across their research sample. Each analysis can be run with a single command and requires no user input beyond the region masks, making the process accessible to a wide range of researchers. The input masks provided by the user are assumed to be in 2 mm isotropic MNI152NLIN6Asym space, as provided with the FSL library (https://fsl.fmrib.ox.ac.uk/fsl/fslwiki). In addition to the PDF report, the tool returns all of the input values and results as a Python object from which the user can extract particular variables for further analysis or for integration in a larger analysis pipeline.

### Gene information

Gene expression information is taken from the Allen Human Brain Atlas microarray dataset (29). This presents mRNA expression values for 3,702 postmortem brain tissue samples obtained from six healthy donors (five male; mean age = 42.5 years, range = 24-57 years). The version of these data used in the tool are previously created cortical representations in MNI152 space, where each image voxel is assigned an imputed value based on the sparsely sampled microarray measurements (32). Reliable expression values are not available for all genes and so, as can be seen in Table 1, some receptor subunits are not represented in the dataset. It should also be noted that the tissue samples are predominantly from the left hemisphere and so right hemisphere values are primarily inferred from the left.

All receptor expression values are normalised relative to the median expression across the cortex (0-1) using a robust sigmoid normalisation method (33). After normalisation, expression values from each 2 mm^3^ image voxel within the input mask are extracted for analysis. Image voxels that do not contain grey matter have no expression estimates and so are excluded. To illustrate, a 30 x 30 x 30 mm^3^ input mask that consists entirely of grey matter would therefore return 3,375 datapoints for analysis. The number of datapoints returned scales with the proportion of grey matter to other tissue types present within the target brain region.

### Excitation/inhibition ratios

An estimated excitation/inhibition (E:I) ratio for each input region is calculated by summing expression values of ionotropic glutamate receptors (NMDA, kainate, and AMPA) within the region mask and dividing by the sum of (ionotropic) GABA_A_ receptors (34).

### Single and Two Region Mapping

The user can input either a single MRS region or two separate regions to be compared. The information output for each option is similar, with the addition of statistical comparison information in the two region case. More specifically, an image showing the location of each mask in MNI space is created first. Relative Glutamatergic and GABAergic subunit expression for each region are extracted using the above methods and plotted using radar and violin plots grouped by receptor type. Normalised expression for neuromodulator receptor genes are also given as radar and violin plots. Radar plots are given in addition to violin plots to aid the user in identifying general patterns in gene expression. An E:I ratio is calculated and included in the output file.

As noted, an additional function in the case of two different regions being input is to statistically compare these. Tables are created alongside the violin plots to statistically compare the central tendencies and distribution shape between the regions. Differences in central tendencies are indicated by percentage difference in means and Cohen’s d. Ninety-five percent confidence intervals for Cohen’s d are estimated through an unbiased bootstrap procedure. Distribution shapes are compared through Kolmogorov-Smirnov tests. D values for Kolmogorov-Smirnov tests range from 0-1, with higher values suggesting greater similarity in the shapes of the distributions and lower values suggesting greater distribution differences.

### Multiple participant voxels

A methodological limitation of MRS in humans is the relatively large volume of the acquisition voxel. This, coupled with challenges in placing voxels consistently, can lead to variations in the tissue sampled across different participants or within individuals across sessions (35). Information about the likely receptor composition of each individual voxel for a study population is therefore useful for understanding how much variability in tissue properties there may be and whether there are any acquisitions that may be particular outliers. The third function of this tool therefore allows for the comparison of two or more voxel masks to explore variability in expression due to minor differences in mask location.

Firstly, a voxel overlap mask is created to demonstrate the consistency in voxel mask placement. This is visualised in the output PDF and is also output as a NIfTI image. For each individual mask, relative expressions will be calculated and plotted using radar and violin plots. The median expression value for each subunit based on all included masks will also be plotted, allowing for a difference from the median score for each individual mask to be calculated as an indicator of variability. Using glutamatergic and GABAergic subunit expressions, an E:I ratio is also calculated for each individual mask. These are both included in the output PDF and output as a separate text file.

### Usage examples

We present three example analyses to illustrate use of the tool. These are: A) A single region analysis based upon an MRS voxel placed in the occipital cortex; B) A two region analysis that contrasts the occipital voxel with one located in the posterior cingulate cortex; and C) An analysis comparing five different participants whose MRS voxels are all located in the medial prefrontal cortex.

## Results

Selected figures from each analysis type are presented here to illustrate the functionality of the tool. Particular observations from the results are noted to highlight the information a user can potentially obtain about their MRS region or regions. Full output PDFs, along with the voxel masks used (in NIfTI format), are available at https://osf.io/rvt2j/.

### Example usage A Single region

To analyse a single MRS region, the user is required to enter the path to the relevant binary mask image. In this example, we point to a mask that covers sections of the occipital and parietal lobes. They may also enter a label for the mask. In this case the label “Occipital” is used. The output PDF will be saved either in the current working directory or in a directory specified by the user. Code to run such and analysis takes the following form:

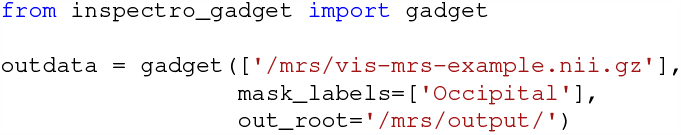

Looking at some selected elements of the output, we can first see a visualisation of the mask in Fig. 1A. The estimated E:I ratio is also shown here. Expression values for each subunit of all available GABAergic, glutamatergic, dopamine, serotonin and adrenergic receptors are presented as series of violin plots, grouped by receptor type or receptor subunit class (Fig. 1B). Zooming in on a single receptor subunit class - GABA_A_ receptor *α* subunits - in Fig. 1C, we are able to note a number of things. Firstly, *α*4 subunits appear to be relatively more likely to be present in this region. At the same time, for the other subunit types we can see what appears to be distributions formed by a mixture of two subdistributions. For example, the GABA_A_ *α*1 distribution has peaks at approximately 0.85 and 0.45. This may reflect the fact that the voxel spans two distinct cortical regions, highlighting the variability in receptor composition across the cortex. Given this potential microstructural variability within the MRS voxel, the researcher may wish to alter its location, depending upon the specifics of their research question.

**Figure 1.**
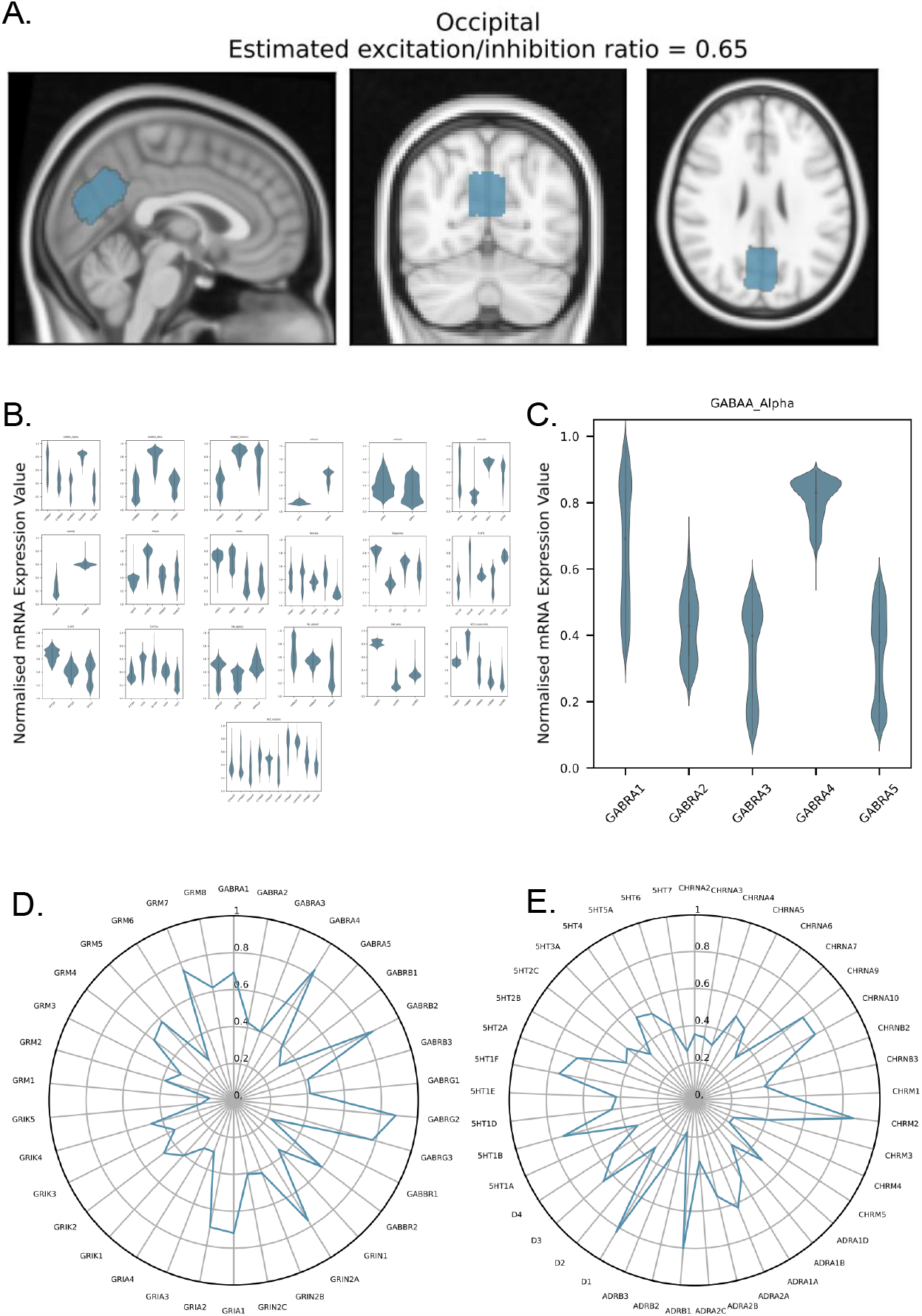
(A) Shows the mask placement for the occipital region of interest in blue in sagittal, axial and radial views. Excitation inhibition ratio within that region is also given. (B) All violin plots produced for the region mask showing normalised mRNA expression values, grouped by receptor type. (C) Violin plot demonstrating normalised mRNA expression value distributions in GABAA *α* subunits. (D&E) Radar plots showing median normalised mRNA expression values for receptor subunits within the region mask.

Normalised expression information is also represented in radar plots, one for excitatory and inhibitory transmitter receptors (Fig. 1D) and one for neuromodulator receptors (Fig. 1E). Looking at Fig. 1D, we can see relatively high expression of GABA_A_ *α*4, *γ*2 and *γ*3 subunits. GABA_B_ receptors appear to have relatively low expression levels in this region. In the case of glutamatergic receptors, we also see a relatively low expression of kainate receptors in the region. On the other hand NMDA2A, AMPA1, AMPA2, and mGlu7 subunits all appear to be relatively highly expressed.

In the case of neuromodulators, shown in Fig. 1E, we can also see significant variability in gene expression across receptor and subunit types. For example, within the serotonergic system there is relatively high expression of 5-HT 1B, 1F, and 2A receptors, but low expression of other 5-HT receptor types. Similarly, we see high expression of D1-type dopaminergic receptors but a relatively lower expression of D2-type ones.

### Example B - Two region comparison

In this second example, the gene expression estimates from two different MRS voxels are compared. The first of these is located in the occipital cortex and the second in the posterior cingulate. To conduct such an analysis, the user enters the path to each mask. They may also enter labels for each region (in this case “Occipital” and “Posterior cingulate”). The output PDF will be saved either in the current working directory or in a directory specified by the user. Code to run such and analysis takes the following form:

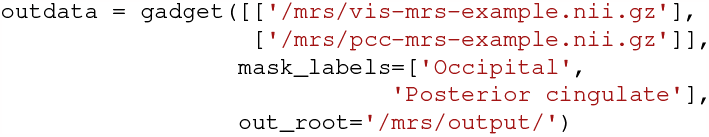

As can be seen in Fig. 2A, images showing each of the voxel locations are created. Matched with these are estimates of the E:I ration for each region. In this case, we can see that the posterior cingulate is estimated to have a higher E:I ratio than the occipital cortex.

**Figure 2.**
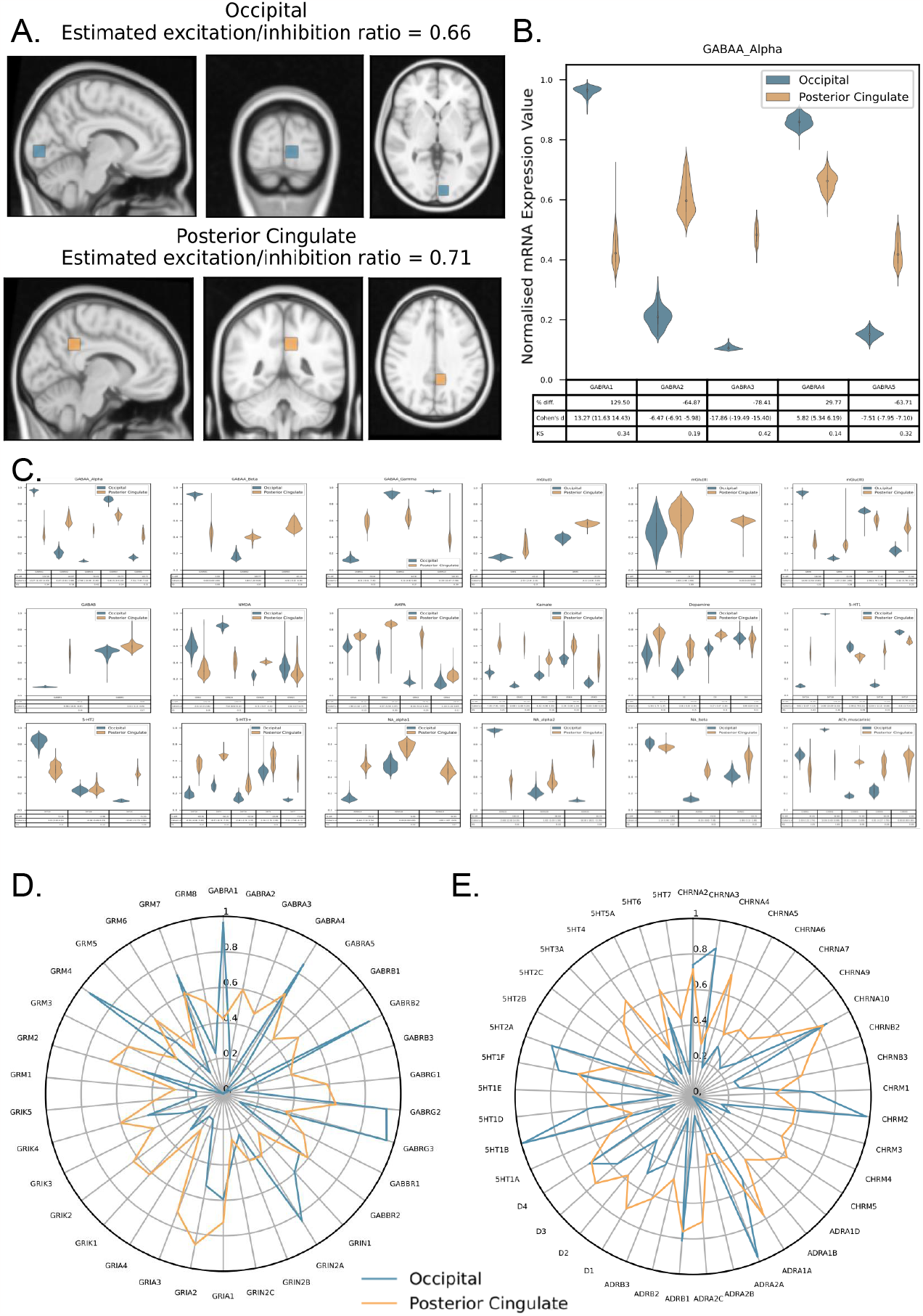
(A) Shows the placement of the voxel masks placed in the occipital lobe in blue and the posterior cingulate in yellow, in sagittal, axial and radial views. Excitation inhibition ratio within that region is also given. (B) Violin plot demonstrating normalised mRNA expression value distributions in GABA_A_ *α* subunits with the output tables below. Blue plots represent the occipital voxel and yellow plots represent the posterior cingulate voxel. (C) All violin plots produced for the region mask showing normalised mRNA expression values, grouped by receptor type. (D&E) Radar plots showing median normalised mRNA expression values for receptor subunits within the occipital voxel (blue) and the posterior cingulate voxel (yellow).

Expression estimates for different genes are then compared between the two regions through side-by-side violin plots and a set of descriptive statistics (Fig. 2B&C). Taking

GABA_A_ receptor *α* subunits as an illustration, clear differences in subunit composition can be seen between the two regions. For example, *α*1 subunits are highly expressed within the occipital cortex but only moderately expressed in the posterior cingulate. This is highlighted by the calculated percentage difference and Cohen’s d statistics (129.5% and 13.27, respectively). In contrast, *α*3 subunits have more similar expression between the two regions, with only a 29.77% difference. The violin plots also allow the distributions of expression values within a voxel to be compared between the regions. From this it can be observed, for example, that *α*1 subunit expression is quite consistent within the occipital voxel but is more varied across the posterior cingulate one. This can be compared to *α*4 expression, which differs in magnitude between both regions but has similar variability in each. Patterns of differences in both expression level and in distribution can also be seen across the other GABAergic, glutamatergic, dopamine, serotonin and adrenergic receptors Fig. 2C.

As with single region analyses, radar plots are produced to give an overview of relative expression across excitatory/inhibitory and neuromodulatory receptors (Fig. 2D&E). Estimates from both input voxels are displayed on the same plot to allow comparison between them. In terms of excitatory/inhibitory receptors, a more even distribution of median expression values can be seen in the posterior cingulate than in the occipital cortex (Fig. 2D). The latter tends more to have higher expression values for a small set of subunits. This may reflect the higher variability in expression values across the input voxel that is often seen for the posterior cingulate voxel. More specific differences for both GABAergic and glutamatergic receptors can also be observed. For example, there is a generally higher expression of kainate receptors in the posterior cingulate, with very low expression of this receptor type in the occipital region. At the same time, GABA_A_ *γ*2 and *γ*3 subunits are highly expressed in the occipital cortex but have low expression estimates in the posterior cingulate. A similar general pattern of more evenly distributed median expression values in the posterior cingulate can also be seen for neuromodulatory receptors (Fig. 2E). In the occipital cortex, high expression of specific receptor types, such as 5-HT1B and alpha2A adrenoceptors can be seen. Muscarinic acetylcholine M1 and M2 receptors are also relatively highly expressed in the occipital cortex. In contrast, these receptor types are not highly expressed in the posterior cingulate but M3, M4 and M5 receptors are (ones that have relatively low expression in the occipital cortex voxel).

### Example usage C - Multiple participants

The final example uses five individual participant anterior cingulate voxel maps to explore variability created by inconsistency in voxel placement between subjects. Paths to each participant mask are entered by the user. The user may also enter participant labels. The output PDF and a voxel overlap NIfTI image are written to the current working directory or to a directory specified by the user. Code to run such and analysis takes the following form:

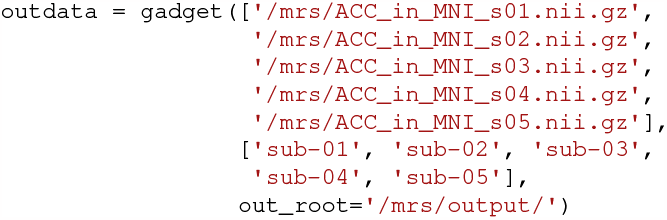

The first element of the multi-subject PDF output is a figure showing the overlap between the participant masks (Fig. 3A). This allows the user to see how consistent voxel placement is in their sample (35). In the example shown, voxel placement is relatively consistent but we can see that at least one participant voxel is located posterior to the others. Next, E:I ratio estimates are shown for each participant, along with the median value for the sample as a whole (Fig. 3B). E:I ratio values are quite consistent across the example sample, although the estimate for “sub-01” is somewhat lower than the others.

**Figure 3.**
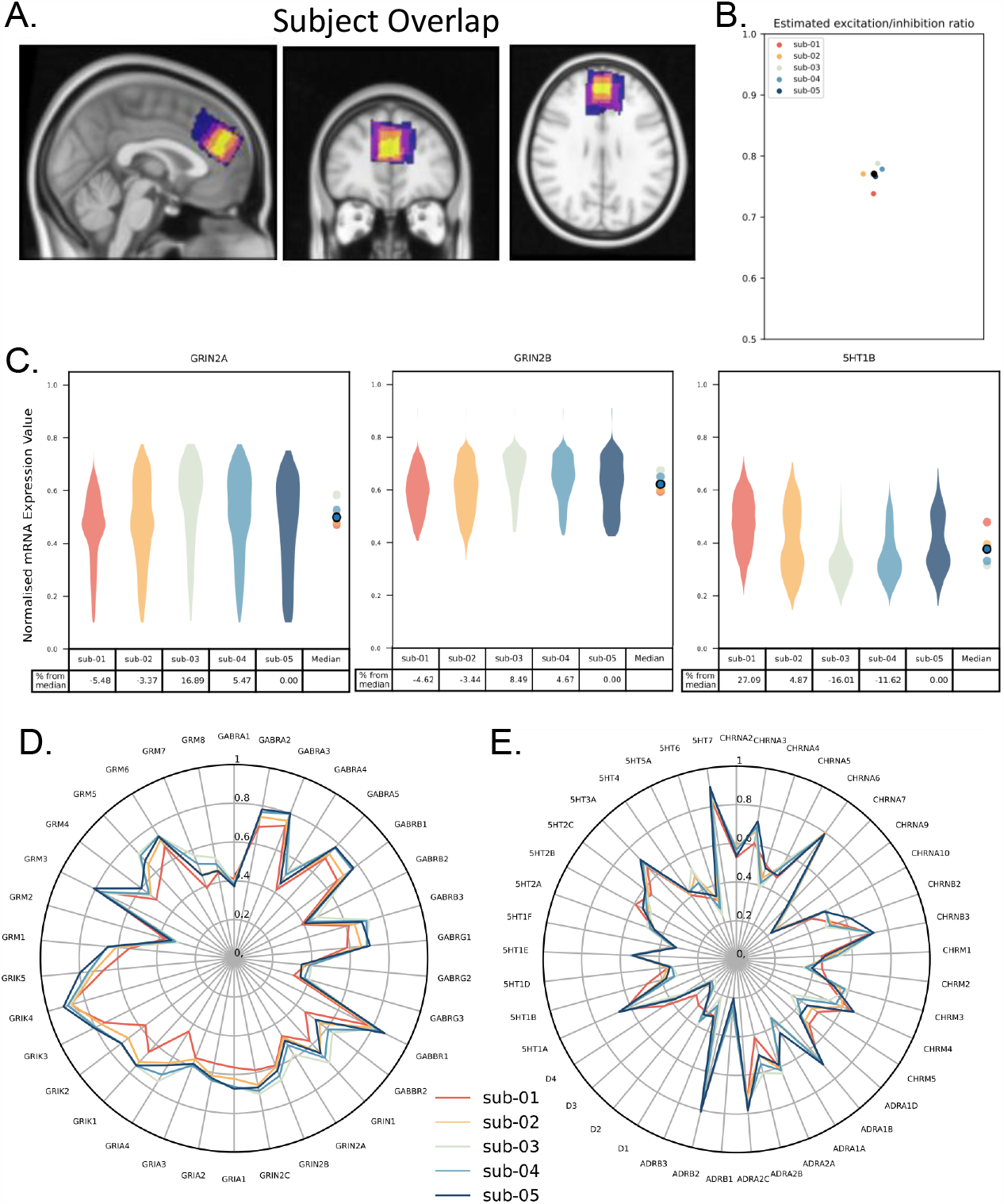
(A) Heat map demonstrating the overlay of medial prefrontal cortex masks used for five subjects (B) Plotted excitation/inhibition ratios for each of the 5 subject region masks. The group median is shown in black. (C) Violin plots produced for the subjects mask showing normalised mRNA expression values for GRIN2A, GRIN2B and 5HT1B subunits. (D&E) Radar plots showing median normalised mRNA expression values for receptor subunits for each subject mask (yellow).

As with the analysis of a single region or comparison between two regions, gene expression estimates are shown via a series of violin plots. In the case of a multi-participant analysis, each gene is shown in a separate plot, with the values for each participant shown (Fig. 3C). In addition to participant violin plots, participant median values are shown as a univariate scatter plot. The sample median is also shown here. How far the expression estimates for a particular participant are from the sample median can be established both through this plot and through percentage deviation values shown in a table below.

Fig. 3C shows these plots for two NMDA receptor subunits (2A and 2B), along with 5-HT1B expression. In the case of NMDA receptor subunit 2B, we can see consistent expression values both within voxels and across all the participants. In contrast, estimates for 2A subunits are more variable within participants (ranging from around 0.1 to 0.8 across the voxel) and one participant, “sub-03”, has a relative expression estimate that is 16.89% higher than the sample median. This may influence how one interprets glutamatergic MRS estimates from this participant.

For 5-HT1B relative expression estimates, some evidence of there being sub-distributions for “sub-02” and “sub-05” can be seen. This may indicate that their voxels are located across a microstructural boundary in the brain. The violin plots for “sub-03” and “sub-04” suggest that their voxel is located mostly to one side of such a boundary, with consistently lower 5-HT1B relative expression estimates. On the other hand, “sub-01” has a median estimate 27.09% higher than the sample median.

Radar plots of relative expression estimates are also produced (Fig. 3D&E). In these, estimates for each participant are plotted to allow a comparison of the general expression landscape between them. Looking at GABAergic and glutamatergic genes in Fig. 3D, “sub-01” stands out as having generally lower relative expression estimates across a number of genes. That participant does not show the same patter of lower expression in the case of neuromodulator genes (Fig. 3E). This contrast between transmitter types may be of particular relevance when considering GABA and glutamte MRS estimates for that participant compared to those making up the rest of the sample.

## Discussion

Here we present ‘InSpectro-Gadget’: A tool that aims to give researchers additional information that they may use to develop a more comprehensive mechanistic understanding of transmitter estimates from MRS data. It does this by giving reference estimates of relative gene expression for a range of relevant neurotransmitter receptor genes. Most importantly, it provides this information in an accessible format with a minimum of user input.

A number of different advantages of the tool are demonstrated in the usage examples above. The first of these is its potential applications in interpreting relationships between MRS measures of neurotransmitters and function. The comparison of the occipital and posterior cingulate cortex voxels revealed a markedly different receptor landscape in each. For example, quite different GABA_A_ *α* subunits can be found in each location. This is of relevance to the link between the transmitter itself (as measured with MRS) and functional outputs as the subunit composition of these receptors is linked, amongst other things, to different effects on postsynaptic cells, to differences in the types of cells present, and to responses to external modulators (36–39). Similarly, the differences in the neuromodulator receptor populations between the two regions will be linked to differences in how those regions respond to changes in behavioural states.

A second potential use of the information obtained is more methodological in nature. From the example analysis of a single region, there was evidence that the MRS voxel lay across the boundary of two different microstructural regions (purposefully introduced in this case). This can also be seen for some participants in the multi-participant example. This information can help the user assess variation in their data and may allow them to modify their voxel site in order to achieve a more homogeneous tissue composition. Taking the multi-participant example, certain participants had potentially outlying relative gene expression patterns compared to the sample as a whole. This information may be of assistance to the user in allowing them to explain potential outlier MRS measures by showing if particular participant voxels are located over an area that has different cytoarchitectural properties to others.

As well as being useful for additional insight into existing MRS datasets, InSpectro-Gadget may also aid in experiment planning. As genetic differences and neurotransmitter systems become increasingly studied, they are likely to become the experimental focus of some MRS studies. For example, one may wish to study relationships between 5-HT2A receptors and GABAergic function (40) or metabolic correlates of NMDA subunit expression in schizophrenia (41). In each case, one may wish to establish optimal target regions for the receptors of interest, both in terms of receptor expression and in practicality of MRS voxel site. To do this, a user may create some test voxels and enter them into InSpectro-Gadget. This will give them a reference map of the transmitter terrain within each potential location. Locations can then be compared for both feasibility (i.e., location in the brain) and appropriateness to the experimental question.

It should be noted that other software is available that interfaces with the Allen Human Brain Atlas. These include the *abagen* (42), *ENIGMA* (43), and *Brain Annotation* (44) toolboxes. Although similar information to that produced by InSpectro-Gadget could potentially be obtained with such alternative tools, it would require extensive input from the user to get the required expression values and then represent them in a usable manner. This difficulty could potentially limit the accessibility of gene expression data to the majority of MRS researchers.

One feature of this tool is to establish the degree to which voxels overlap within a group of participants. The relevance of such variability in voxel localisation has been widely discussed in the MRS literature and is well recognised amongst users (35, 45, 46). InSpectro-Gadget provides additional information to users by allowing them to identify potentially relevant variation in the microstructural context they are measuring from across participants. This may allow them to make more informed decisions about excluding certain participants. It may also contribute to the interpretation of variability in data obtained from voxels that are not perfectly overlapping with each other.

## Limitations

While this tool can provide valuable insights into the relative subunit expression within region masks, caution should be taken when using this to interpret meaning and generalise findings. The gene information used for this tool comes from the Allen Human Brain Atlas microarray dataset, which uses only a very small and potentially unrepresentative sample of human brains (29). This dataset includes detailed gene expression data from six human healthy adult brains, predominantly males aged 24-57 and as such, may not be fully representative of sex, age or ethnicity based gene expression diversity in the brain. Brain morphology and subunit expressions have been shown to vary between sexes (47), and change across the lifespan (48), thus highlighting why a lack of diversity in gene information may be limiting to interpretations. Additionally, the cortical representation of the Allen human brain expression data used in this study may be more accurately representative of expression in the left hemisphere than the right given that of the eight hemisphere samples included, only two were right hemisphere. While steps were taken to limit the impact of this when creating the interpolated whole brain maps, the authors did note that this would limit investigation of brain lateralization effects with this data (32).

## Conclusions

InSpectro-Gadget is an easy to use tool that aims to give neuroimaging researchers an additional layer of information to integrate in their research. The tool may be of use to a wide range of users from both a methodological and interpretive perspective. Inherent limitations of the gene data must be noted, but these may highlight the utility of future development of individualised measures of multiple transmitter systems in the human brain.

## Acknowledgements

This work was supported by grants from the Taiwan Ministry of Science and Technology to NWD (110-2628-H-038-001-MY4) and by the Taiwan Ministry of Education Higher Education Sprout Project. This preprint was created using a modified version of Ricardo Henriques’ BioRxiv LaTex template (https://henriqueslab.github.io/resources/bioRxivTemplate/).

## Data availability

Code is available at https://github.com/lizmcmanus/Inspectro-Gadget. Data for reproducing the analyses described here are available at https://osf.io/rvt2j/.

## Author contributions

EM: Conceptualization, Formal Analysis, Methodology, Visualization, Writing – original draft;

NM: Conceptualization, Writing – original draft;

NWD: Conceptualization, Formal Analysis, Methodology, Visualization, Writing – original draft

